# Deciphering the genetic structure of the Quebec founder population using genealogies

**DOI:** 10.1101/2022.09.19.508491

**Authors:** Laurence Gagnon, Claudia Moreau, Catherine Laprise, Hélène Vézina, Simon L. Girard

**Affiliations:** Département des sciences fondamentales, Université du Québec à Chicoutimi, Saguenay, Québec, G7H 2B1, Canada; Centre Intersectoriel en Santé Durable (CISD), Université du Québec à Chicoutimi, Saguenay, Québec, G7H 2B1, Canada; Centre Intégré Universitaire en Santé et Services Sociaux du Saguenay–Lac-Saint-Jean, Saguenay, Québec, QC G7H 7K9, Canada; Département des sciences humaines et sociales, Université du Québec à Chicoutimi, Saguenay, Québec, G7H 2B1, Canada; Projet BALSAC, Université du Québec à Chicoutimi, Saguenay, Québec, G7H 2B1, Canada; Centre de recherche CERVO, Université Laval, Québec, Québec, G1V 0A6, Canada

## Abstract

Using genealogy to study the demographic history of a population makes it possible to overcome the models and assumptions often used in population genetics. The Quebec founder population is one of the few populations in the World having access to the complete genealogy of the last 400 years. The goal of this paper is to follow the evolution of the Quebec population structure generation per generation from the beginning of European colonization until the present day. To do so, we calculated the kinship coefficients of all ancestors’ pairs in the ascending genealogy of 665 individuals from eight regional and ethnocultural groups per 25-year period. We show that the Quebec population structure appeared in the St. Lawrence valley as early as 1750. At that time, the ancestors of two groups, the Sagueneans and the Acadians from the Gaspé Peninsula, experienced a marked increase in kinship and inbreeding levels which have shaped the contemporary population structure. Interestingly, this structure arose before the colonization of the Saguenay region and at the very beginning of the Gaspé Peninsula settlement. The resulting regional founder effects in these two groups, but also in the other regional groups, led to differences in the present-day identity-by-descent sharing and are directly linked to the number of most recent common ancestors and their genetic contribution to the studied subjects.

## Introduction

Founder populations have been particularly helpful in demonstrating how past demographic events have shaped present-day genetic structure and its consequences on human health (1–3). Studying the past demographic history of a population often relies on current genetic data and models of ascending genealogical trees (4,5). However, developing efficient methods for inferring the underlying genealogy has proved challenging (6,7) or requires lots of contemporary and ancient genomes data (8). To avoid using such assumptions, one would need the complete genealogy of the population. Few populations in the World have access to such genealogical data (9–11). The Quebec province of Canada relies on the BALSAC population register, a large collection of linked data from parish records, to reconstruct the genealogy of the vast majority of Quebecers, mostly of French Canadian descent, but also of other origins, since the foundation of the colony in the 17^th^ century until recent times (12). This invaluable resource allows the detailed mapping of the population structure over time. Indeed, it has been shown that the genealogical lines covering the last 400 years explain most of the present-day genetic structure of the Quebec population (13,14).

This paper will focus on Quebecers genealogically anchored into five regions (from west to east): the Montreal and Quebec City areas, the Saguenay-Lac-St-Jean (Saguenay) and North Shore regions, and the Gaspé Peninsula (Gaspé) where four subgroups were sampled (Acadians, French Canadians, Loyalists, and Channel Islanders). Most Quebecers of French Canadian ancestry are descendants of around 8,500 settlers who came predominantly from France between 1608 and 1760 (15). These European newcomers first settled in Quebec City (1608) and Montreal (1642) which are now two major urban regions (Figure 1A) and along the shores of the St.Lawrence river. Following the British Conquest of 1760, French immigration decreased dramatically, and the French-speaking population expanded mostly through natural increase. Population growth led to the colonization of new regions, including more remote and isolated regions, favoring population subdivision (13).

**Figure 1.**
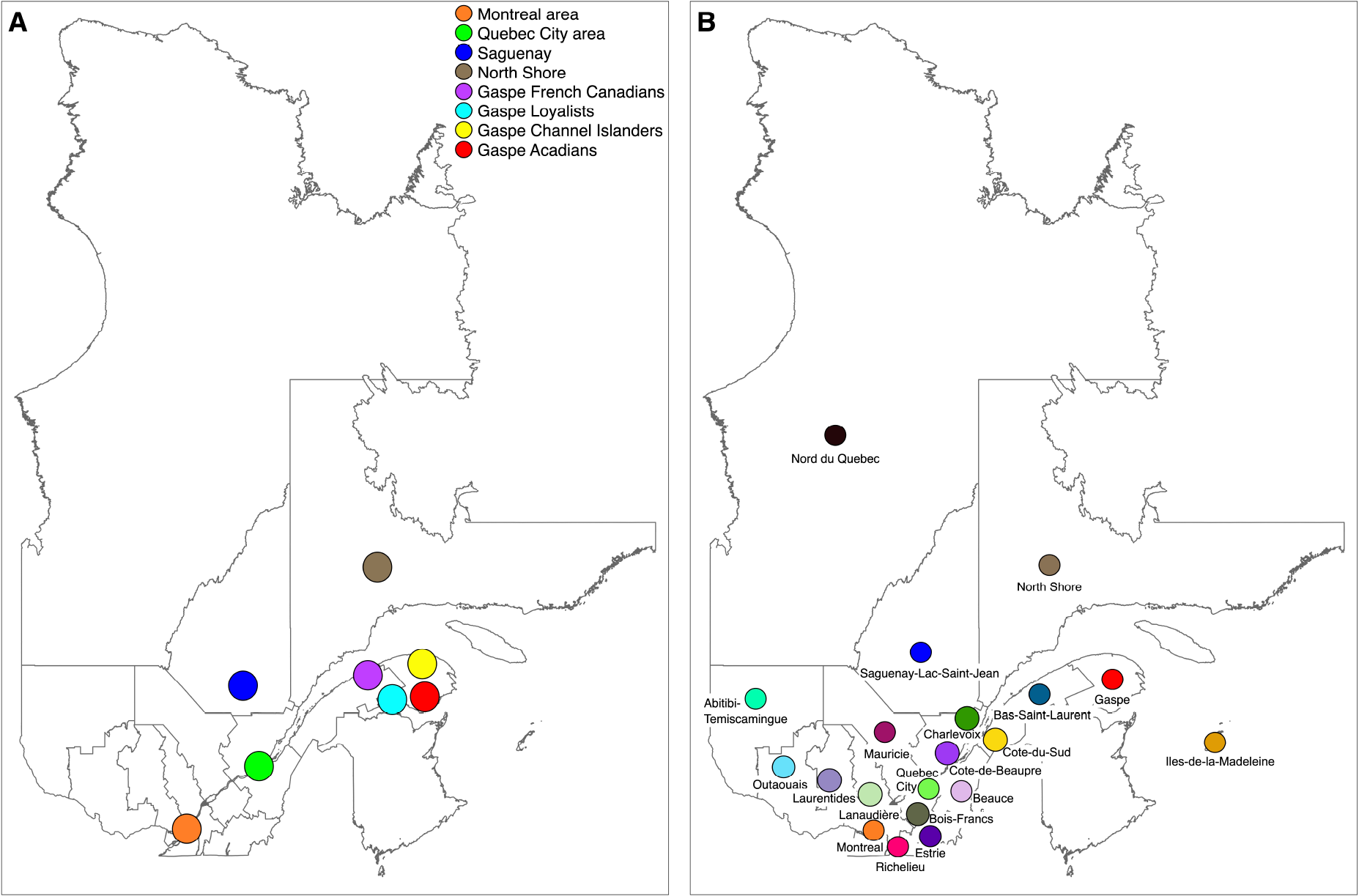
The geographical location of groups investigated in this study (A) and parents’ regions of marriage that are used in figure 2 (B) Genetic data is often used to study the contemporary population structure (20) and we have previously shown that the genetic structure of the Quebec population is well correlated with the one inferred using genealogical measures (13,21,22). However, genealogies are an invaluable tool to study how the population structure was shaped in the past generations. The goal of this paper was to follow the evolution, generation per generation, of the Quebec regional population structure from its colonization until the present day. To do so, we looked at the kinship and inbreeding levels of all ancestors in the genealogies. We also tried to explain the population fine-structure inferred with present-day genetic identity-by-descent (IBD) sharing using genealogical measures.

Permanent European settlement in Gaspe began during the second half of the 18^th^ century with the arrival of Acadians, who escaped deportation by the British (16). They were soon joined by English-speaking United Empire Loyalists who chose to remain under British rule after the American Declaration of Independence in 1776. From 1830-1840, many Quebecers of French Canadian ancestry from the lower part of the St. Lawrence valley also settled in the Gaspé peninsula (16). At the same time, a fourth group, inhabitants of the Channel Islands, came to Gaspé for the fishing industry.

The settlement of Saguenay started in 1838 with founders mostly coming from the neighboring region of Charlevoix which was colonized earlier by the end of the 17^th^ century. The Saguenay population size underwent a 25-fold increase between 1861 and 1961, mostly due to a high birth rate (17,18), while the whole Quebec population increased only 5-fold. The western part of the North Shore was colonized by ancestors who came from the Charlevoix and Bas-St-Laurent regions (19) while the eastern part pioneers were mostly fishermen from Iles-de-la-Madeleine and Gaspe.

## Subjects and Methods

This study was approved by the University of Quebec in Chicoutimi (UQAC) ethics board. Written informed consent was obtained from all adult participants.

### Cohort

The data consist of 579 individuals from the Quebec Regional Reference Sample and 86 unaffected individuals from the Saguenay-Lac-St-Jean asthma familial cohort (Figure 1A and Table 1)(13,21,23). The individuals are distributed in five regions and eight groups (based on geographical and ethnocultural criteria) of the province of Quebec: the Montreal and Quebec City areas, the Saguenay and North Shore regions, and the Gaspé Peninsula. For the latter, four subgroups were sampled, namely Acadians, French Canadians, Loyalists, and Channel Islanders. Individuals were sampled regardless of their proportion in the population. To ensure regional connection, individuals needed to have their 4 grandparents born in the Quebec province and 1 or 2 parents born in the particular region except for the Montreal area where only first criteria (the four grandparents) was applied. The four ethnocultural subgroups of Gaspé were self-reported. A strong correspondence between ancestral origins traced in genealogies and self-reported origins was found in a previous study (24). Genetic and genealogical data are available for the 665 individuals.

**Table 1.** Regional and ethnocultural groups’ description.

| **Group** | **Abbreviation** | **Sample Size** | **Average year of Parents' Marriage** |
| --- | --- | --- | --- |
| Montreal Area | MTL | 138 | 1950 |
| Quebec City Area | QUE | 70 | 1945 |
| Saguenay-Lac-Saint-Jean | SAG | 86 | 1977 |
| North Shore | NSH | 47 | 1949 |
| Gaspé French Canadians | GFC | 97 | 1945 |
| Gaspé Loyalists | GLO | 71 | 1939 |
| Gaspé Channel Islanders | GCI | 67 | 1941 |
| Gaspé Acadians | GAC | 89 | 1942 |

### Genealogical Data and Analyses

Genealogical data were obtained through the BALSAC project (12). Genealogies were reconstructed with average completeness of at least 60% up to the tenth generation for all groups, except for the Loyalists and Channel Islanders of Gaspé (explained in part by their later time of arrival in Quebec), consistent with previous results (13)(Figure S1). The completeness is the proportion of ancestors found at each generation in the genealogy compared to the maximum possible number of ancestors. Information on the parents’ year (±5 years for confidentiality concerns) and region of marriage or if outside Quebec, country of origin was obtained for 94,076 distinct individuals throughout the genealogy. There are 20 regions of marriage in our data (Figure 1B). Parents’ marriage years were grouped into 25-year periods to minimize the parent-child overlap within the same period, the average time between parents’ marriage and their children’s marriages in our data being 32 years.

The kinship coefficient at the maximum generational depth was computed using the R GENLIB library v1.1.6 (22) for each pair of ancestors within 25-year periods. Multidimensional scaling (MDS) was performed on the pairwise kinship distance matrix (1-kinship coefficient). Ancestors who had no kinship ties with anybody were removed from this analysis. The average pairwise kinship and inbreeding coefficients for the ancestors of each group were calculated for each period. Genealogical inbreeding of an individual is equal to the kinship coefficient of his/her parents. To avoid group size bias on the mean ancestors’ kinship and inbreeding, we randomly resampled 47 individuals (the smallest group size (Table 1)) 1,000 times.

Most recent common ancestors (MRCAs) were counted for the subjects’ pairs within groups for each distance in meioses (for example, there is four meioses between two cousins). The expected genetic contribution (CG), consisting in summing the transmission probabilities over all genealogical paths connecting an ancestor to a descendant assuming that parents transmit half of their genome to each child, was also calculated for each MRCA to both descendants using GENLIB. The GC product to both individuals was summed over all MRCAs. Groups were resampled down to 47 subjects 1,000 times again to avoid size bias.

### Genetic Data and Analyses

Genotyping was conducted on Illumina Omni Express (700,000 SNPs) and Illumina Omni 2.5 chips (2.5M SNPs) chips. Both chips have been merged to keep only common SNPs. Quality control filters were applied at the individual and SNP levels using PLINK software v1.9 (25). We retained individuals with at least 98% genotypes among all SNPs. At the SNP level, we retained SNPs with at least 98% genotypes among all individuals, located on the autosomes and in Hardy–Weinberg equilibrium p > 0.001, yielding 663,172 SNPs. Closely related individuals (first cousins, kinship coefficient >= 0.0625) were eliminated, yielding a final sample size of 665 individuals (Table 1). A PCA was performed using PLINK software to confirm that our final dataset reflects the previously described Quebec population structure (13,21) (Figure S2).

The assessment of pairwise IBD segments was performed using refinedIBD software v17Jan20 (26). This software was selected for its power and accuracy in detecting IBD segments.

## Results

### How Population Structure was Shaped

The evolution of the Quebec population structure was assessed using pairwise kinship coefficients of all ancestors at the maximum depth for each period (Figure 2). In this figure, we used the parents’ marriage region (Figure 1B) to color each dot (representing an ancestor whose parents married in that period). Before 1750, the ancestors from three regions (Côte-de-Beaupré, Côte-du-Sud, and Charlevoix), clustered together and are differentiated from another group of immigrants, whose parents did not marry in Quebec but whose country of origin is known (see interactive Figure 2 for countries of origin). This group is composed of ancestors mainly coming from Acadia (83%) and a lower proportion from France (5%). The present-day population structure appears as early as 1751-1775 distinguishing Charlevoix and the ancestors of the Gaspé Acadians (GAC). By 1826-1850, the first Saguenean ancestors appeared and the population structure at that time was very similar to the one depicted on the PCA of the present-day subjects (Figure S2).

**Figure 2.**
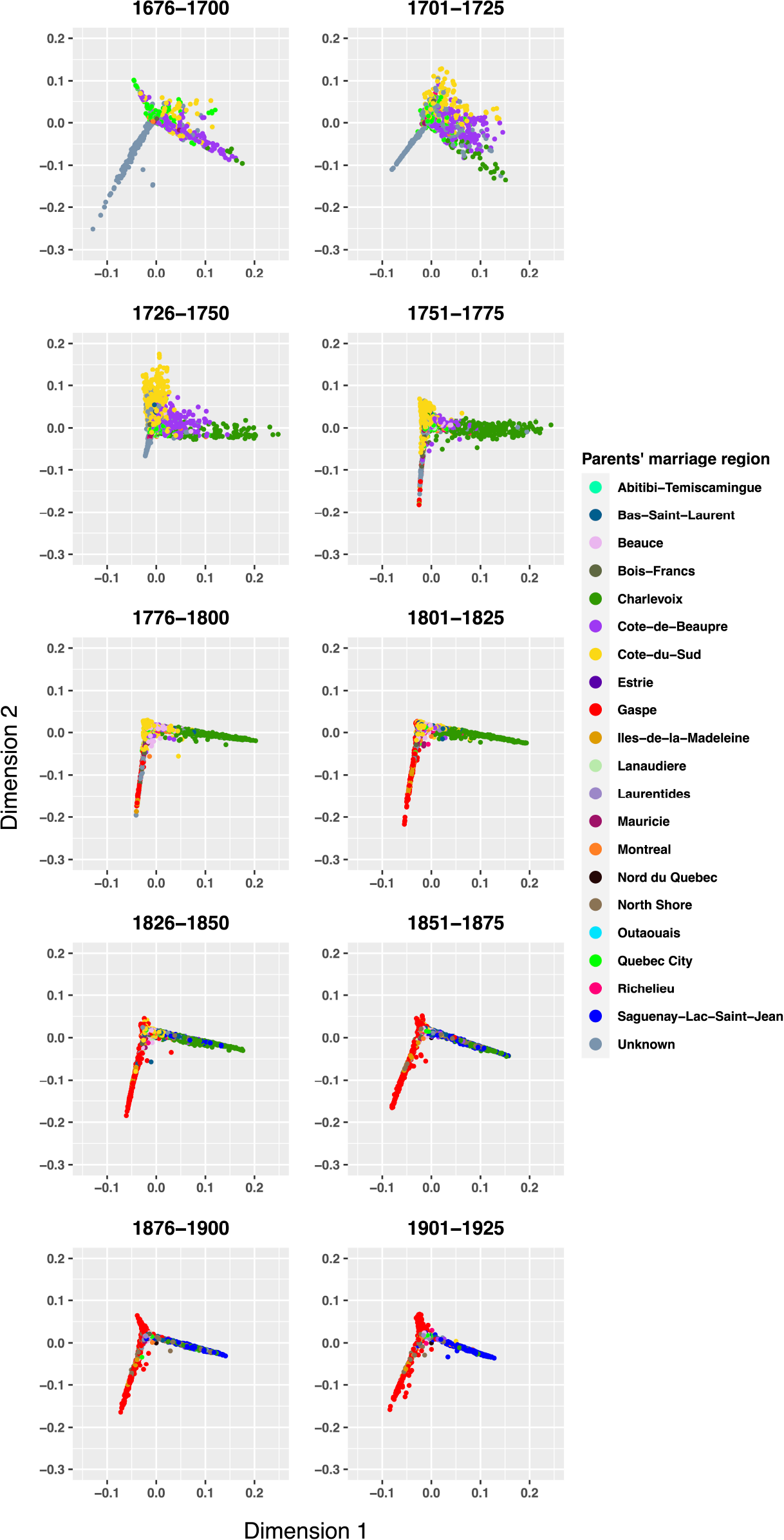
Multidimensional scaling (MDS) of the pairwise kinship coefficients of ancestors per 25-year period. MDS was performed on the individuals’ pairwise kinship distance matrix, (i.e., 1-kinship coefficient) of ancestors whose parents were married at each period. The pairwise kinship coefficient was computed using the R GENLIB library. The external interactive version of this figure is available at https://laugag17.github.io/quebec_founder_pop_interactive_figure/figure2_interactive_graph.html.

### Mean Kinship and Inbreeding Over Time

We averaged for each group the genealogical kinship and inbreeding coefficients at the maximum depth for ancestors living at each period (based on parents’ marriage date) (Figure 3A and S3A). From 1750, the GAC and Saguenay ancestors went through a marked increase in averaged kinship compared to the other groups. By 1825, the GAC ancestors had continued to increase while the Saguenay ancestors had reached a plateau. The ancestors’ mean kinship increase in GAC and Saguenay groups was obviously accompanied by an increase in inbreeding (Figure 3B and S3B). Until 1850, the average inbreeding coefficient at the maximum depth was higher for the Saguenay ancestors. After 1850, the GAC ancestors exceeded the Saguenay ancestors, whereas the latter reached a plateau. The three other Gaspé groups’ average inbreeding coefficient increase was slower at the beginning, but almost reached the Saguenay ancestors’ inbreeding value by 1925. Interestingly, for all groups the increase in inbreeding was more important than the increase in kinship levels.

**Figure 3.**
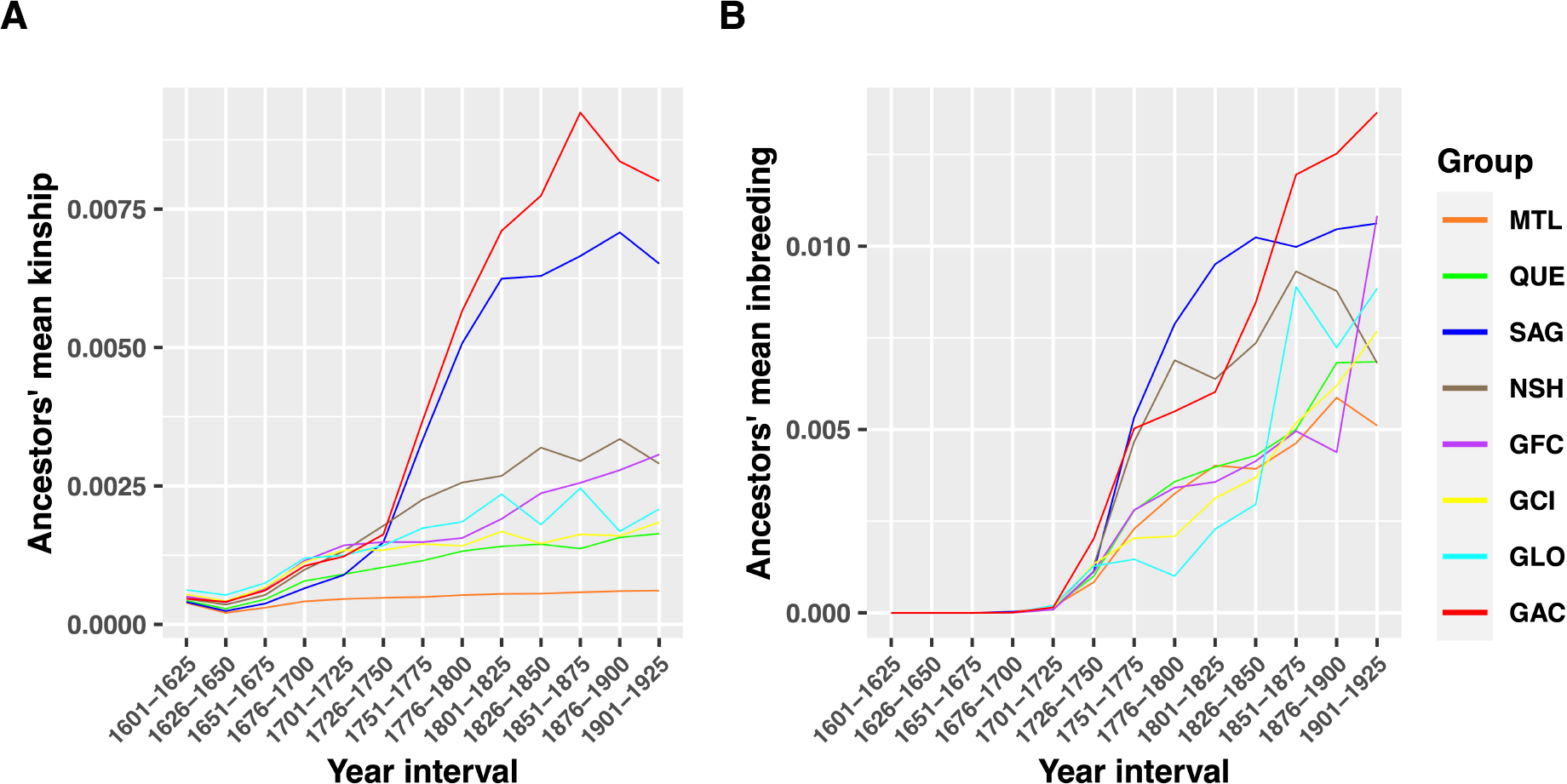
Average kinship (A) and inbreeding (B) coefficients within groups per 25-year period. The solid line represents the mean values of the 1,000 bootstraps of 47 individuals.

### IBD Sharing and most Recent Common Ancestors

For each group, we plotted the mean number of IBD segments shared among subjects’ pairs per segment length (bins of 5cM) and compared this with the cumulative MRCA counts per meiosis (Figure 4A and 4B). Note that MRCAs are not unique so the same MRCAs can appear for many subjects’ pairs and they will be counted each time they appear. These two metrics, one using genetic data and the other using genealogical data, show very similar patterns. Interestingly, the Sagueneans’ pairs shared more IBD segments of short lengths (<25cM), but less of long lengths (>25cM) compared to the North Shore and the four Gaspé groups leaving only the urban and older groups (Montreal and Quebec City areas) behind for longer segments. This is also reflected by the cumulative MRCA count until 10 meioses (Table S1). Inversely, the Gaspé Loyalists (GLO) shared less short segments and more long segments. Note that the MRCA count for the GLO is biased towards the right of the graph (Figure 4B) since their completeness decreases faster than the other groups (Figure S1). This was also observed for the Gaspé Channel Islanders to a lesser extent.

**Figure 4.**
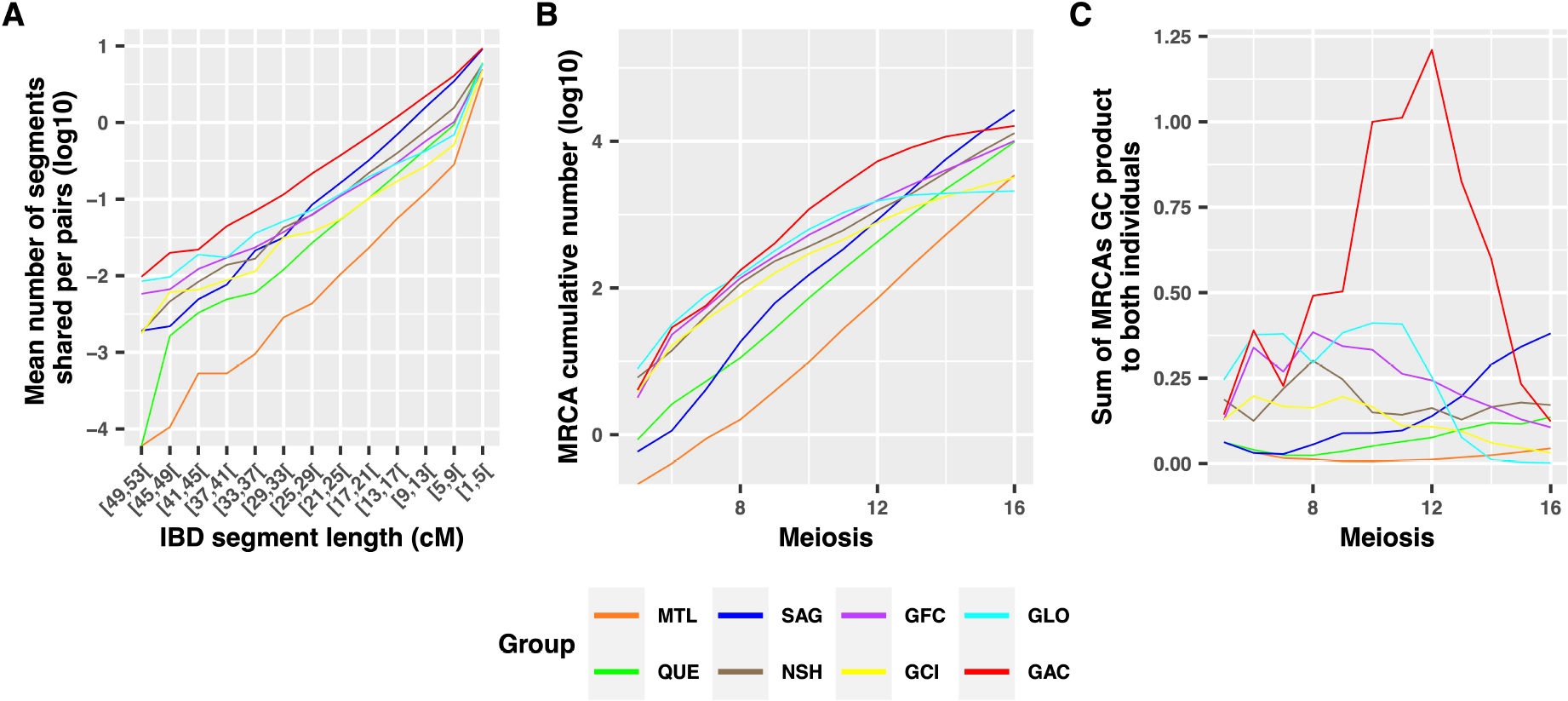
Within groups IBD segment sharing by length (A) as well as MRCA cumulative count (B) and genetic contribution (C) per meiosis. IBD segment lengths are binned into 5cM intervals. The MRCA cumulative count (B) and the sum of genetic contribution (C) presented in this figure are the averages of 1,000 bootstraps of 47 individuals.

We also calculated in the genealogies the product of the genetic contributions (GC) of each MRCA to both individuals and then we summed these products for all MRCAs. Figure 4C presents this GC sum averaged for 1,000 bootstraps of 47 individuals (see also Figure S4 for intervals). Note the very high GC of GAC MRCAs starting at a few meioses from now. For the three other Gaspé groups and the North Shore, closer MRCAs also had a higher GC than those of Montreal, Quebec City, and Saguenay. However, for the latter, the GC sum is higher for more distant MRCAs.

## Discussion

In this study, we applied an innovative method to show how the Quebec founder population structure was shaped over time. We found that the previously described structure differentiating the GAC and the Saguenay groups (13) emerged early in the colonization process (1750), almost a hundred years before the colonization of the Saguenay region (1840) (27–29) (Figure 2). At this time, Saguenay ancestors were mostly located in the Charlevoix region where they had established only two or three generations before. The small number of founding families in Charlevoix followed by a rapid expansion in Saguenay in the 19^th^ century led to changes in the frequencies of alleles and diseases (30). Similarly, GAC descend from a few founding families, but they did not go through a rapid expansion. Instead, they mostly married inside their community due to linguistic and cultural barriers present with the other Gaspe groups (31). Additionally, they were the only ones in the area until 1780 (16). Both groups’ colonization started with a few founding families implicating a smaller number of founders which can explain the increase in average kinship levels. Despite their different subsequent colonization processes, Saguenay and GAC ancestors have a very similar mean kinship increase (Figure 3A) starting in 1750 when the contemporary population structure appeared. For both Saguenay and GAC ancestors, spouses had higher chances of being related than those in the other groups. However, the average inbreeding was higher among Saguenay ancestors until 1850 when GAC ancestors went through a marked increase which lasted until recent times. This is consistent with previous findings showing that close inbreeding is the lowest in the province for Sagueneans and the highest for Gaspe (28) (Figure S5). In fact, for both kinship and inbreeding, the Saguenay ancestors reached a plateau at the time the region was colonized (1838) and the expansion started (around 1860) while GAC never went through such a rapid expansion. This increase in inbreeding followed by stabilization among Saguenay ancestors was previously explained by the evolution of nonrandom mating as well as by the evolution of inbreeding resulting from drift (32). This would need further investigation to understand its implications on the contemporary population. Nevertheless, the GAC and the other Gaspe ethno cultural groups did not reach such a plateau.

The regional fine structure could be observed within groups by comparing IBD sharing patterns (Figure 4A). The GAC and Saguenay groups share more IBD segments <25cM than the other groups, in line with previous results (21) and consistent with their higher mean kinship. However, for long segments >25cM, the IBD sharing of Sagueneans’ pairs decreases more rapidly than the Gaspé and North shore groups. This is explained by the recent MRCA counts from 5 to 10 meioses (Figure 4B and Table S1) which are more numerous in the Gaspé and North Shore groups than in the Sagueneans. In other words, the Sagueneans have less recent, but more distant common ancestors than other eastern groups. This is consistent with the close inbreeding being less important in Saguenay than among the Gaspé and the North Shore subjects (Figure S5) (28). As MRCA can appear many times in this analysis and even though the Saguenean subjects descend from fewer founders, they have more numerous MRCAs (after 13 meioses), which means that some of them appear very often, consistent with an expanding population due to a high birth rate and also observed in previous studies (27). Another interesting ethnocultural group is the Gaspé Loyalists (GLO) which is genetically different from other Quebec groups on the third PC (Figure S2). The GLO is among the groups with the lowest mean number of short IBD segments shared per pair, but they reach the second highest mean number of pairs sharing IBD (after the GAC group) above 25cM, consistent with previous results (21). Indeed, GLO ancestors did not undergo an early marked kinship increase like Saguenay and GAC ancestors, they only show an inbreeding increase after 1850 (Figure 3B) which is consistent with the highest inbreeding values found at the 6-7 generations for GLO subjects (Figure S5). The GLO group comes from more numerous and diversified founders which could have affected the sharing of short segments (33). After their settlement, the GLO ancestors have remained quite isolated for more than 150 years as shown by their rapid inbreeding increase (Figure 3B), which could have exacerbated their sharing of long IBD segments. This is again consistent with previous findings (31) and explains the particular IBD sharing among GLO subjects.

In this paper, we show a direct relation between the IBD segment sharing and the cumulative number of genealogical MRCAs per pair (Figure 4A and 4B). Shared IBD segment length distribution depends on the number of common ancestors and the distance connecting both individuals to their common ancestor (22). The chance of transmitting a segment also depends on the GC of the common ancestor to both descendants (22). Thus, common ancestors who contributed a lot to the present-day gene pool would be more susceptible to transmitting an IBD segment than those who contributed less. Usually, the closer the ancestors are to their descendants, the bigger their GC is (Figure 4C). But in Saguenay, there are unusually great contributors among distant ancestors (27). The number of MRCAs above 20 meioses is in the same order of magnitude for Saguenay as for Montreal and Quebec City individuals. However, the Saguenay ancestors’ GC above 10 meioses was higher and the resulting IBD sharing of shorter segments (less than 25cM) is also higher. Sagueneans share more segments of less than 25cM than any other group except GAC. In turn, GAC close common ancestors have a larger GC and they have the highest IBD sharing for all length bins even if their close MRCA counts (until 8 meioses) are similar to the other Gaspé groups and the North Shore subjects.

Some limitations are present in this work. The genealogical completeness is not consistent across all groups (Figure S1) but was left uncorrected to retain two groups of Gaspé that would have been filtered out otherwise (Gaspé Loyalists and Channel Islanders) (13,21). This explains the aberrant curves in figure 4B, especially for Loyalists’ MRCA counts above 12 meioses. We also reported an inconsistent number of participants across groups (Table 1). To overcome this, a bootstrap method was performed for specific genealogical analyses (Figures 3, 4B, and 4C). Finally, a generation gap was present between the Saguenay and the other groups (Table 1) that was not accounted in figure 4 since the IBD sharing to be compared with genealogical MRCAs also includes this generation gap. For the other genealogical analyses presented in this study, this was not relevant since ancestors of each period were grouped regardless of the subjects’ generation.

In conclusion, genealogies are an invaluable tool to study the evolution of the population structure over time and to understand how the present-day genetic structure was shaped. We have shown that the Quebec population structure subdividing the Saguenay and GAC groups appeared early in the history of the province, even before the colonization of the Saguenay region. At that time, they both experienced a marked average kinship and inbreeding increase until recent times, when Saguenay reached a plateau. The resulting strong founder effect that occurred led to differences in the present-day IBD sharing and is directly linked to less numerous recent, but more numerous distant MRCAs for the Sagueneans compared to the GAC. Another understudied group, the GLO, was shown to have numerous recent MRCAs resulting in higher sharing of long IBD segments compared to all other groups except GAC. However, as their founders were more numerous and diversified and also due to their lower close MRCA GC, they did not go through a kinship increase during the same period as the Saguenay and GAC groups and their resulting founder effect is less striking.

## Acknowledgments

This work was supported by funding from the CIHR (#420021) in the SLG lab. It was also made possible by the Digital Research Alliance of Canada which provided access to storage and computing resources. We are extremely grateful to all participants in this research. We would like to thank Damian Labuda and his team for the Quebec Regional Reference Sample cohort constitution. LG received funding from the FRQS, the FQRNT, and the CIHR. CL is the director of the CISD (http://www.uqac.ca/santedurable)and the chairholder of the Canada Research Chair in the Environment and Genetics of Respiratory Diseases and Allergy (http://www.chairs.gc.ca).

## Author Contributions

All authors acquired data and approved the final version of the manuscript. LG and CM played an important role in interpreting the results. LG, CM and SLG conceived and designed the study and drafted the manuscript. CL and HV revised the manuscript.

## Data and Code Availability

The 665 individuals’ genealogies and genotypes are available upon request to BALSAC (12). The code used for this study can be found in the following GitHub repository: https://github.com/laugag17/quebec_founder_pop.

## Supplemental Information Legends

**Figure S1. Groups’ mean completeness per generation**

The completeness is the proportion of ancestors present in the genealogy at each generation compared to the maximum possible number of ancestors.

**Figure S2. Principal component analysis of genotype data**

**Figure S3. Average kinship (A) and inbreeding (B) coefficients within groups per 25-year period**

The solid line represents the mean and the dashed lines are the maximum and minimum values of 1,000 bootstraps of 47 individuals.

**Figure S4. MRCA cumulative count (A) and genetic contribution (B) per meiosis within groups**

The solid line represents the mean and the dashed lines are the maximum and minimum values of 1,000 bootstraps of 47 individuals.

**Figure S5. Mean inbreeding coefficient of subjects per group per generation**

The inbreeding coefficient depends on the number of common ancestors present in both parents’ genealogy. Close inbreeding (until 4 generations) provides information on the choice of a spouse while distant inbreeding rather reflects the demographic history of the population.

## References

1. Kere J. Human population genetics: lessons from Finland. Annu Rev Genomics Hum Genet. 2001;2:103–28.

2. Scriver CR. Human Genetics : Lessons from Quebec Populations. Annual Review of Genomics and Human Genetics. 2001;2:69–101.

3. Locke AE, Steinberg KM, Chiang CWK, Service SK, Havulinna AS, Stell L, et al. Exome sequencing of Finnish isolates enhances rare-variant association power. Nature. 2019;572(7769):323–328.

4. Marchi N, Schlichta F, Excoffier L. Demographic inference. Current Biology. 2021;31(6):R276–R279.

5. Tournebize R, Chu G, Moorjani P. Reconstructing the history of founder events using genome-wide patterns of allele sharing across individuals. PLOS Genetics. 2022 juin;18(6):e1010243.

6. Rasmussen MD, Hubisz MJ, Gronau I, Siepel A. Genome-Wide Inference of Ancestral Recombination Graphs. PLoS Genetics. 2014;10(5):e1004342.

7. McVean GAT, Cardin NJ. Approximating the coalescence with recombination. Philosophical Transactions of the Royal Society B: Biological Sciences. 2005;360(1459):1387–1393.

8. Wohns AW, Wong Y, Jeffery B, Akbari A, Mallick S, Pinhasi R, et al. A unified genealogy of modern and ancient genomes. Science. 2022 Feb 25;375(6583):eabi8264.

9. Helgason A, Hrafnkelsson B, Gulcher JR, Ward R, Stefánsson K. A Populationwide Coalescent Analysis of Icelandic Matrilineal and Patrilineal Genealogies: Evidence for a Faster Evolutionary Rate of mtDNA Lineages than Y Chromosomes. The American Journal of Human Genetics. 2003;72:1370–1388.

10. Pluzhnikov A, Nolan DK, Tan Z, McPeek MS, Ober C. Correlation of intergenerational family sizes suggests a genetic component of reproductive fitness. American Journal of Human Genetics. 2007;81(1):165–169.

11. Pettay JE, Kruuk LEB, Jokela J, Lummaa V. Heritability and genetic constraints of life-history trait evolution in preindustrial humans. Proceedings of the National Academy of Sciences of the United States of America. 2005;102(8):2838–2843.

12. BALSAC. BALSAC [Internet]. BALSAC. Available from: https://balsac.uqac.ca/.

13. Roy-Gagnon MH, Moreau C, Bherer C, St-Onge P, Sinnett D, Laprise C, et al. Genomic and genealogical investigation of the French Canadian founder population structure. Human Genetics. 2011 May;129(5):521–31.

14. Anderson-Trocmé L, Nelson D, Zabad S, Diaz-Papkovich A, Baya N, Touvier M, et al. On the Genes, Genealogies, and Geographies of Quebec [Internet]. bioRxiv; 2022 [cited 2022 Aug 8]. p. 2022.07.20.500680. Available from: https://www.biorxiv.org/content/ 10.1101/2022.07.20.500680v1.

15. Charbonneau H, Desjardins B, Légaré J, Denis H. The population of the St-Lawrence Valley, 1608-1760. In: A Population History of North America. 2000:99–142.

16. Desjardins M, Frenette Y, Bélanger J. Histoire de la Gaspésie. Éditions de l’IQRC. Revue d’histoire de l’Amérique française. 1999. 797 p.

17. Pouyez C, Lavoie Y. Les Saguenayens. Introduction à l’histoire des populations du Saguenay. Presse de l’UQ. 1983. 386 p.

18. Jette R, Gauvreau D, Guérin M. Aux origines d’une région: le peuplement fondateur de Charlevoix avant 1850. Histoire d’un génome Population et génétique dans l’est du Québec Québec: Presses de l’Université du Québec. 1991;75–106.

19. Frenette P. Histoire de la Côte-Nord. Sainte-Foy, Québec: Institut québécois de recherche sur la culture; 1996. (Collection Les régions du Québec). 667 p.

20. Nait Saada J, Kalantzis G, Shyr D, Cooper F, Robinson M, Gusev A, et al. Identity-by-descent detection across 487,409 British samples reveals fine-scale population structure and ultra-rare variant associations. Nat Commun. 2020 Nov30;11(1):6130.

21. Gauvin H, Moreau C, Lefebvre JF, Laprise C, Vézina H, Labuda D, et al. Genome-wide patterns of identity-by-descent sharing in the French Canadian founder population. Eur J Hum Genet. 2014 Jun;22(6):814–21.

22. Gauvin H, Lefebvre JF, Moreau C, Lavoie EM, Labuda D, Vézina H, et al. GENLIB: an R package for the analysis of genealogical data. BMC Bioinformatics. 2015 May 15;16(1):160.

23. Laprise C. The Saguenay-Lac-Saint-Jean asthma familial collection: the genetics of asthma in a young founder population. Genes Immun. 2014 Apr;15(4):247–55.

24. Vézina H, Tremblay M, Lavoie ÈM, Labuda D, Stringer L. Concordance Between Reported Ethnic Origins and Ancestral Origins of Gaspé Peninsula Residents. Population (English Edition, 2002-). 2014;69(1):7–27.

25. Purcell S, Neale B, Todd-Brown K, Thomas L, Ferreira MAR, Bender D, et al. PLINK: A Tool Set for Whole-Genome Association and Population-Based Linkage Analyses. Am J Hum Genet. 2007 Sep;81(3):559–75.

26. Browning BL, Browning SR. Improving the accuracy and efficiency of identity-by-descent detection in population data. Genetics. 2013 Jun;194(2):459–71.

27. Bherer C, Labuda D, Roy-Gagnon MH, Houde L, Tremblay M, Vézina H. Admixed ancestry and stratification of Quebec regional populations. American Journal of Physical Anthropology. 2011;144 : 432–441.

28. Vézina H, Tremblay M, Houde L. Mesures de l’apparentement biologique au Saguenay-Lac-St-Jean (Québec, Canada) à partir de reconstitutions généalogiques. Annales de démographiehistorique. 2004;108:67–83.

29. Lavoie EM, Tremblay M, Houde L, Vézina H. Demogenetic study of three populations within a region with strong founder effects. Community Genet. 2005;8(3):152–60.

30. Bouchard G, De Braekeleer M. Histoire d’un genome: Population et genetique dans l’est du Quebec. Sillery, Québec: Presses de l’Université du Québec; 1991. 607 p.

31. Moreau C, Vézina H, Yotova V, Hamon R, Knijff PD, Sinnett D, et al. Genetic heterogeneity in regional populations of Quebec - Parental lineages in the Gaspe Peninsula. American Journal of Physical Anthropology. 2009 Aug;139(4):512–22.

32. Mourali-Chebil S, Heyer E. Evolution of inbreeding coefficients and effective size in the population of Saguenay Lac-St.-Jean (Quebec). Hum Biol. 2006 Aug;78(4):495–508.

33. Matthews GJ, Gentilcore RL. Historical Atlas of Canada [Internet]. University of Toronto Press; 1993 [cited 2022 Aug 1]. Available from: http://www.jstor.org/stable/10.3138/9781442675759.

