## Supplementary Material for "Deciphering the genetic structure of the Quebec founder population using genealogies"

**Supplemental Information**

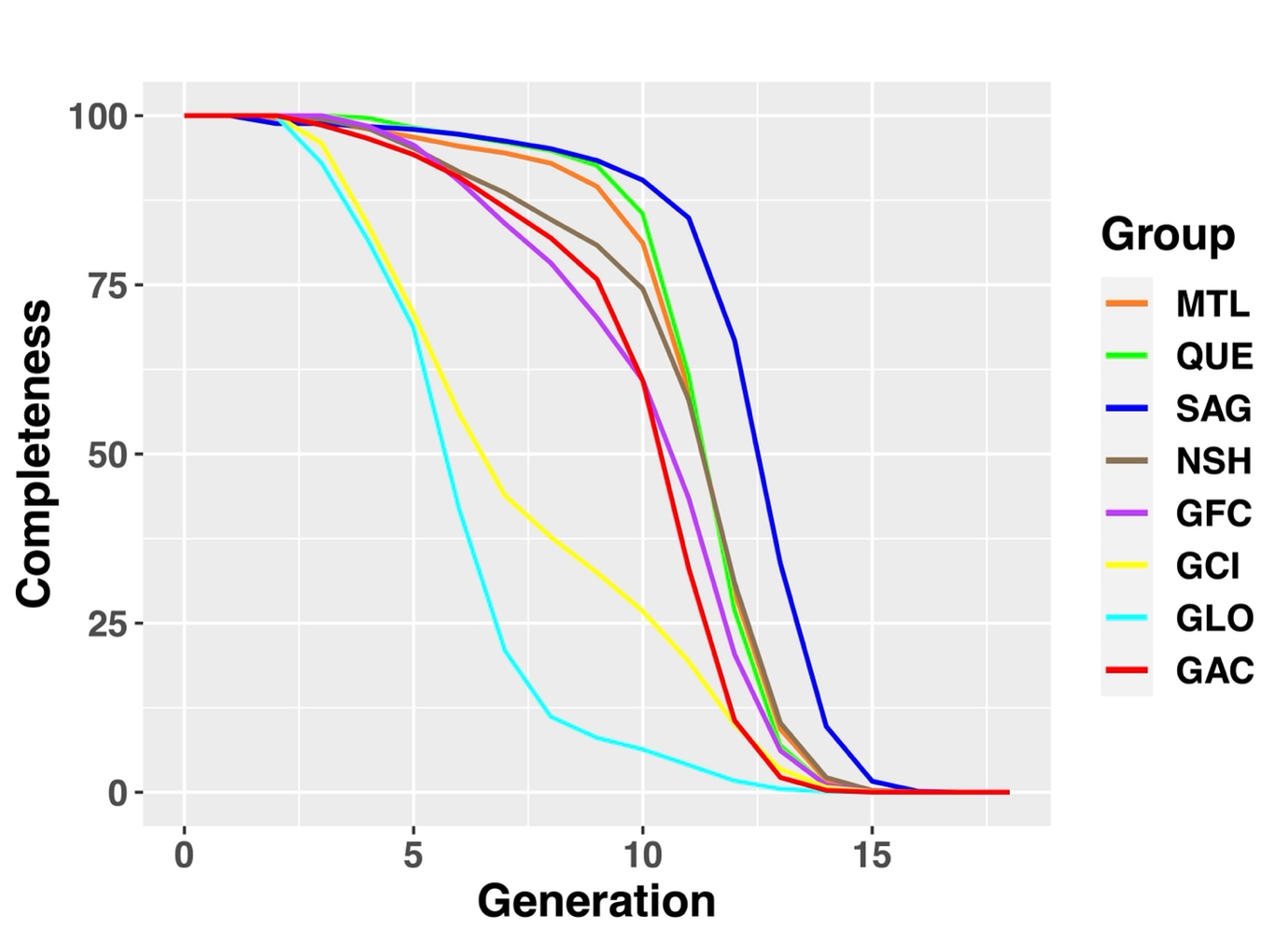

**Figure S1. Groups’ mean completeness per generation**

The completeness is the proportion of ancestors present in the genealogy at each generation compared to the maximum possible number of ancestors.

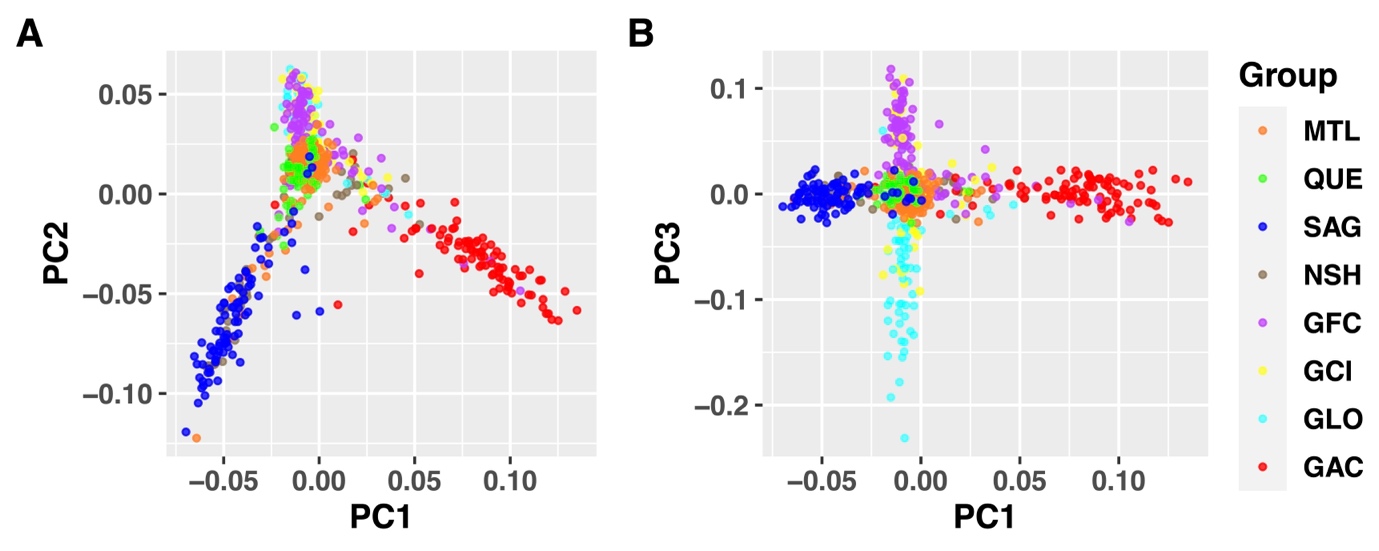

**Figure S2. Principal component analysis of genotype data**

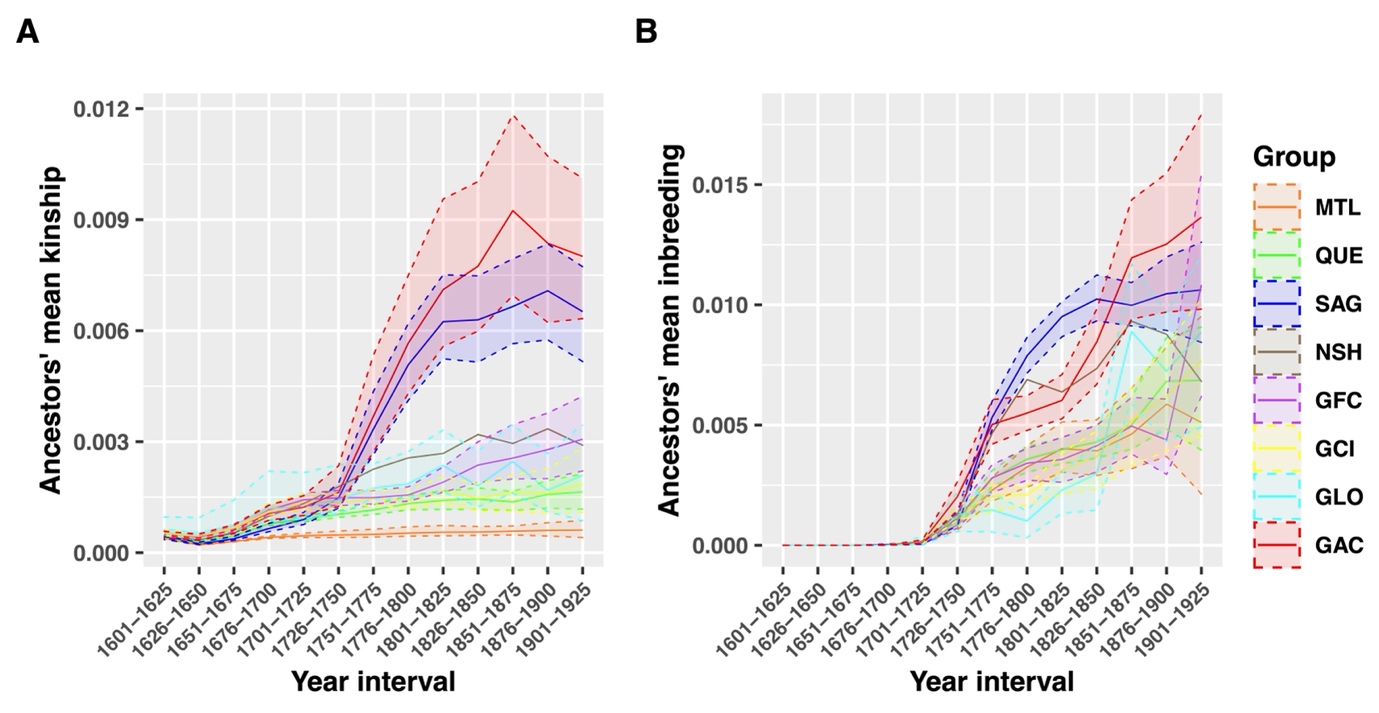

**Figure S3. Average kinship (A) and inbreeding (B) coefficients within groups per 25-year period**

The solid line represents the mean and the dashed lines are the maximum and minimum values of 1,000 bootstraps of 47 individuals.

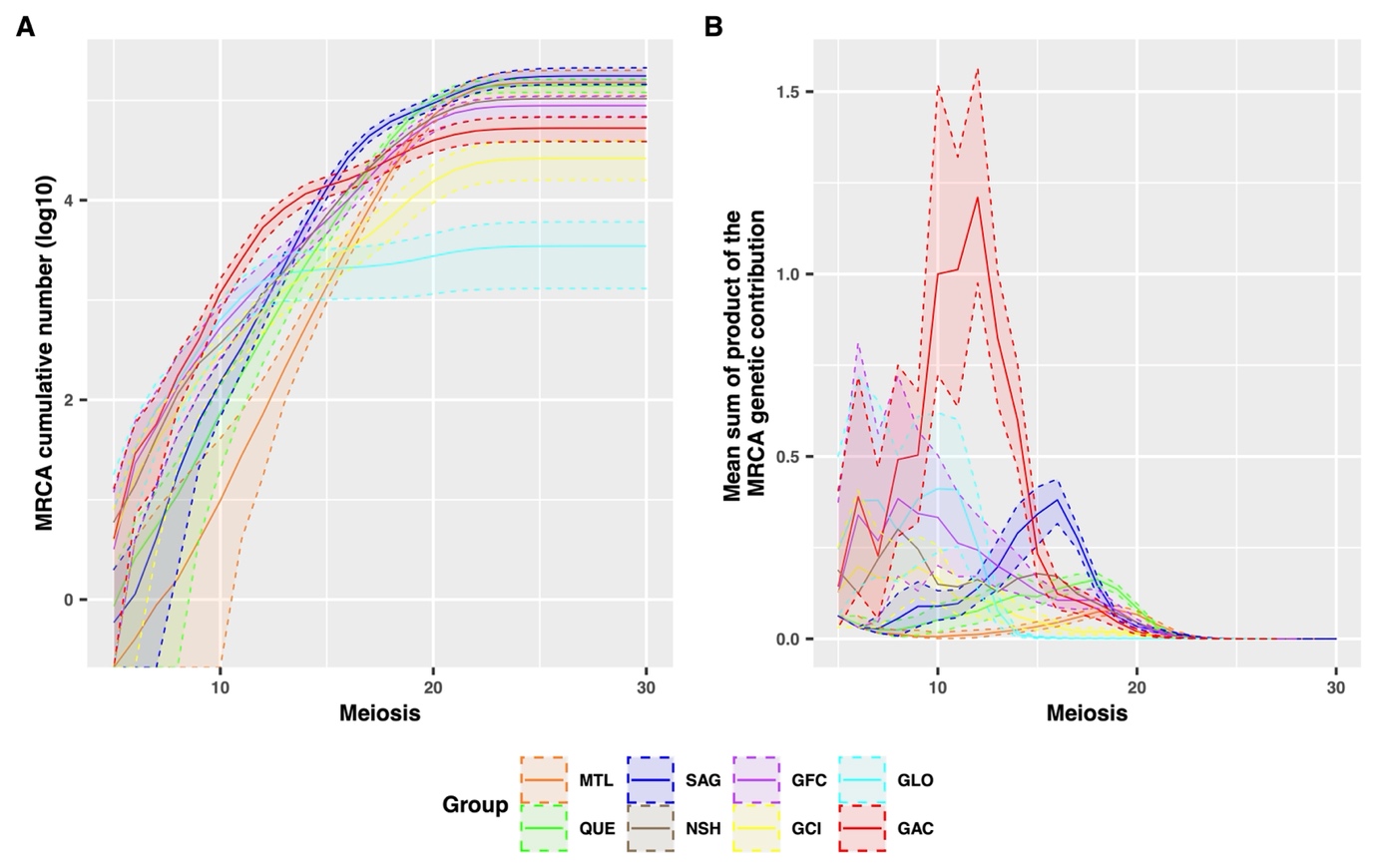

**Figure S4. MRCA cumulative count (A) and genetic contribution (B) per meiosis within groups**

The solid line represents the mean and the dashed lines are the maximum and minimum values of 1,000 bootstraps of 47 individuals.

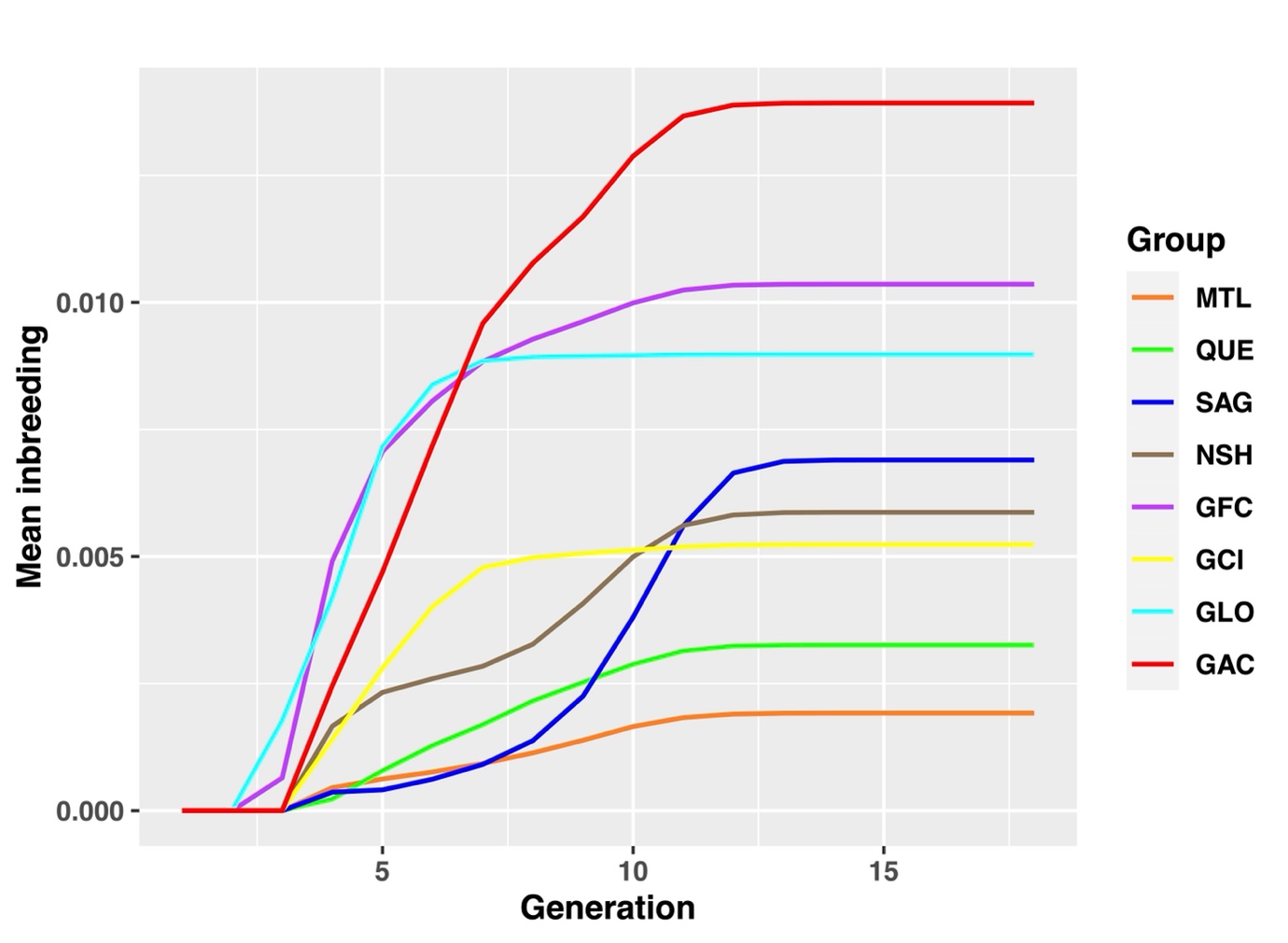

**Figure S5. Mean inbreeding coefficient of subjects per group per generation**

The inbreeding coefficient depends on the number of common ancestors present in both parents' genealogy. Close inbreeding (until 4 generations) provides information on the choice of a spouse while distant inbreeding rather reflects the demographic history of the population.

**Table S1. Cumulative MRCA count**

Mean of 1,000 bootstraps of 47 individuals.

|  | **6** | **8** | **10** | **12** | **14** | **16** | **18** | **20** | **22** | **24** | **26** | **28** | **30** |
| --- | --- | --- | --- | --- | --- | --- | --- | --- | --- | --- | --- | --- | --- |
| MTL | 0 | 2 | 10 | 72 | 534 | 3441 | 19239 | 70845 | 129160 | 148650 | 151048 | 151164 | 151165 |
| QUE | 3 | 11 | 73 | 428 | 2238 | 9779 | 40217 | 100058 | 134125 | 139772 | 140152 | 140154 | 140154 |
| SAG | 1 | 18 | 151 | 845 | 5701 | 26733 | 61315 | 93452 | 139673 | 168269 | 175136 | 175891 | 175902 |
| NSH | 14 | 115 | 367 | 1142 | 3756 | 12913 | 33438 | 67604 | 95445 | 103129 | 104070 | 104097 | 104097 |
| GFC | 23 | 136 | 529 | 1559 | 4048 | 10120 | 28829 | 61358 | 82731 | 87996 | 88586 | 88612 | 88612 |
| GCI | 17 | 77 | 292 | 783 | 1767 | 3214 | 6734 | 15264 | 23194 | 25801 | 26118 | 26119 | 26119 |
| GLO | 32 | 150 | 630 | 1535 | 1954 | 2100 | 2281 | 2740 | 3239 | 3430 | 3462 | 3463 | 3463 |
| GAC | 29 | 175 | 1181 | 5322 | 11577 | 16182 | 25855 | 39748 | 49457 | 52188 | 52614 | 52621 | 52621 |
